# FlashPack: Fast and simple preparation of ultra-high performance capillary columns for LC-MS

**DOI:** 10.1101/426676

**Authors:** Sergey Kovalchuk, Ole N. Jensen, Adelina Rogowska-Wrzesinska

## Abstract

Capillary ultra-high-pressure liquid chromatography (cUHPLC) is essential for in-depth characterization of complex biomolecule mixtures by LC-MS. We developed a simple and fast method called *FlashPack* for custom packing of capillary columns of 50-100 cm length with sub-2-μm sorbent particles. *FlashPack* uses high sorbent concentrations of 500-1000 mg/ml for packing at relatively low pressure of 100 bar. Column blocking by sorbent aggregation is avoided during the packing of sorbent particles by gentle mechanical tapping of the capillary proximal end by a slowly rotating magnet bar. Utilizing a standard 100 bar pressure bomb, *Flashpack* allows for production of 15-25 cm cUHPLC columns within a few minutes and of 50 cm cUHPLC columns in less than an hour. Columns exhibit excellent reproducibility of back-pressure, retention time and resolution (CV 8,7 %). *FlashPack* cUHPLC columns are inexpensive, robust and deliver performance comparable to commercially available cUHPLC columns. The FlashPack method is versatile and enables production of cUHPLC columns using a variety of sorbent materials.

## Introduction

Capillary liquid chromatography (LC) is the central analytical separation technique for liquid chromatography-mass spectrometry (LC-MS) based functional proteomics in biology, biomedicine and clinical medicine. Combined with optimized and often multi-dimensional sample prefractionation, LC-MS approach allows very detailed characterization of the human proteome. However, prefractionation comes at the cost of extended analysis time due to the need for LC-MS processing of each individual fraction [1]. The effectiveness of LC-MS itself can be improved by the application of ultra-high performance capillary chromatography (cUHPLC). For example, analyte separation using a 50 cm length capillary column packed with 2 μm sorbent particles allowed profiling of the yeast proteome (~5,000 proteins) in just about one hour [2]. Further development of ultra-high-performance and high-sensitivity LC separations is very important [3], but is hampered by the fact that only few types of commercial cUHPLC columns are available, the choices of sorbents are limited, and commercial capillary columns come at high costs. A method for efficient, fast and simple custom preparation of 50-100 cm capillary columns for robust and reproducible cUHPLC applications could change this situation.

The most popular setup for capillary column packing today consists of a container (a 2 ml vial) with a stirred sorbent suspension placed into a special “pressure bomb”, which is connected to a nitrogen gas tank (Fig. 1a) [4]. Upon pressurization the sorbent suspension is squeezed into the capillary open end dipped into the suspension. The sorbent is trapped and retained by a frit in the distal capillary end, i.e. a glass frit [5] or a self-assembling sorbent frit in a tapered column end [6].

**Fig. 1.**
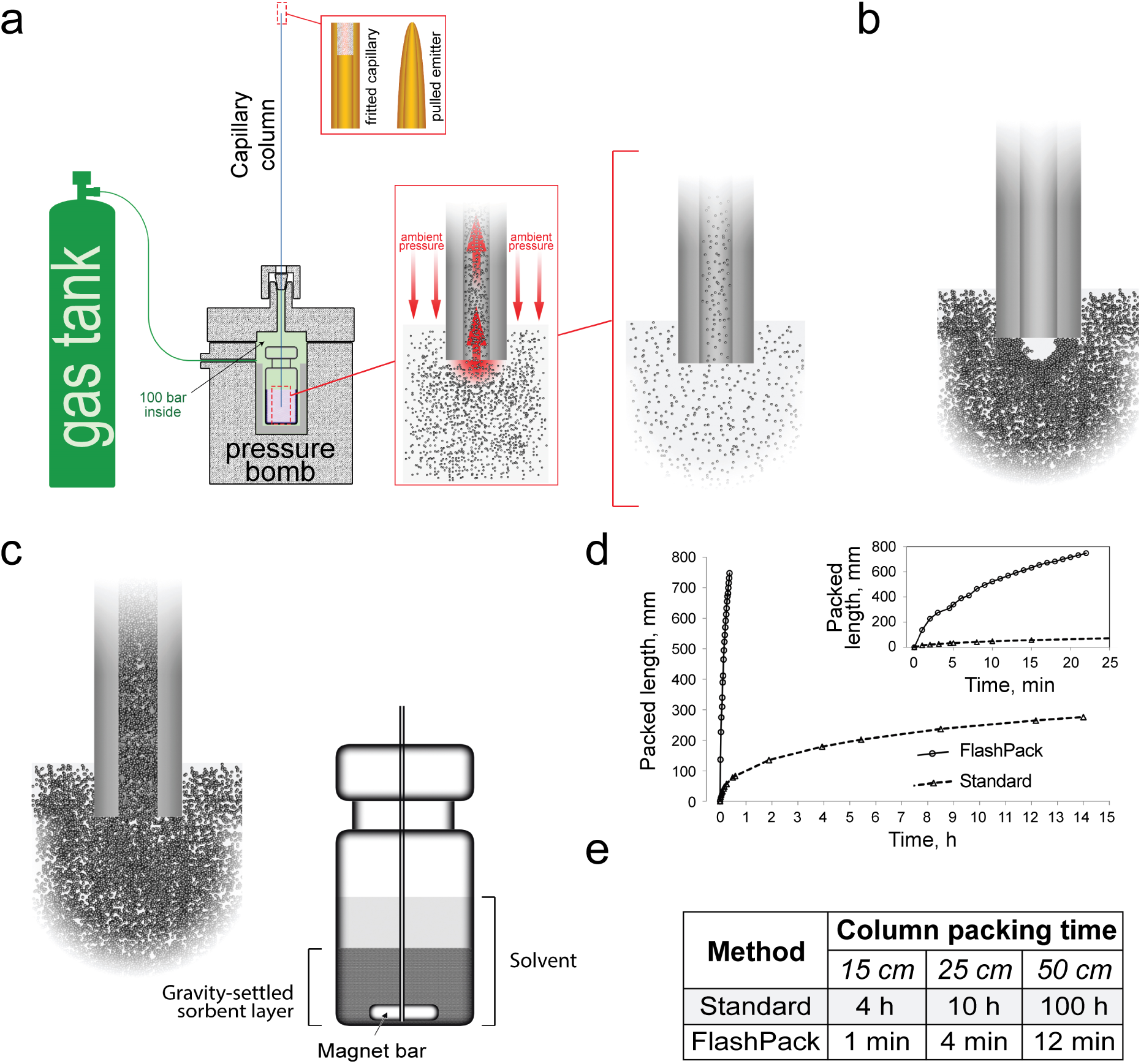
Capillary chromatography column packing using the *FlashPack* protocol at low pressure and high sorbent concentration. **(a)** Schematic illustration of capillary column packing method using a pressure bomb. **(b)** Low sorbent concentration allows only very slow packing rates. High sorbent concentration presumably leads to cupola formation and blocking of the capillary column entrance that disrupts column packing. **(c)** The *FlashPack* method uses mechanical destabilization of the sorbent cupola at the proximal capillary end and allows for very fast packing of capillary columns at ultra-high sorbent suspension concentration and low pressure (100 bar). Mechanical destabilization is carried out by a rotating magnet bar in contact with the silica capillary inside the sorbent vial. **(d)** Column packing time increases exponentially with column length. The FlashPack method reduced packing time dramatically so that a capillary column of 50 cm length can be packed in less than an hour **(e)**.

Capillary column packing is usually accomplished by *“low pressure”* packing using a 100 bar pressure bomb and “low *sorbent concentration*” (2.5-25 mg/ml) [7-9]. The approach is very popular due to its simplicity and is widely used in proteomics LC-MS laboratories. However, the *low pressure/low concentration* method is extremely time-consuming and practically unsuitable for packing of very long (50-100 cm) capillary UHPLC columns (Supplementary Fig. S1). Efficient cUHPLC column packing can be achieved at *“high pressures"* of 1000-4100 bar with *“high sorbent concentrations"* (100-250 mg/ml). This approach is technically more demanding due to the ultra-high pressure conditions [10-12].

We hypothesized that a simple and fast packing method can be developed using the *low pressure/high sorbent concentration* combination provided that clogging and blocking of the column entrance by sorbent aggregation can be avoided during the packing process (Supplementary Fig. S1a). We developed an optimized *FlashPack* approach, which uses a standard pressure bomb at 100 bar to achieve ~100-times higher column packing rates than before, and approaches the efficiency of ultra-high pressure packing protocols. The *FlashPack* protocol is versatile and simple to implement and the produced cUHPLC columns exhibit performance comparable to that of similar commercially available columns.

## Experimental procedures

### Materials

All chemicals and solvents were of LC gradient or LC-MS grade (Sigma); Polyimide coated fused silica capillary (360 μm OD, 30-200 μm ID) was from PostNova. Chromatographic sorbent Inertsil ODS3 2 μm (GLSciences) was used in all LC-MS experiments shown here; other tested chromatography resins included Inertsil ODS3 3 μm (GLSciences), Reprosil Pur C18AQ 1.9, 3 and 5 μm (Dr. Maisch), BEH C18 1.7 μm (Waters), Aeris Peptide 2.6 μm and 1.7 μm, Luna 2 C18 3 μm (Phenomenex), Zorbax SB-C18 1.8 μm (Agilent), Triart C18 1.9 um (YMC), polyLC and polyCAT (polyLC Inc.). The EASY-Spray™ PepMap RSLC C18 2 μm (50 cm*75 μm) column was obtained from Thermo Scientific. The pressure bomb was from Proxeon, Denmark. Similar pressure bomb devices (pressure injection cell, capillary packing unit) are also available from Next Advance and Nanobaume.

### Pulled-emitter (taper-tip) and fritted capillary preparation

Precut and polished fused silica capillaries were flushed with methanol and dried with N_2_ flow. Emitters for taper-tip columns were pulled with a P2000 laser puller (Sutter, USA) according to the manufacturer’s instructions. ESI emitters were visually inspected under a microscope. ESI needle tip diameter for 75 μm ID/360 mm OD fused silica capillary was ~5 μm (proportionally smaller and larger for other capillary IDs). Integrated frits for fritted capillary columns were made using glass-paper (GF/C, Whatman) soaked in Kasil-formamide mixture [13].

### Capillary column packing (general procedure)

Columns were packed into prepared fused silica fritted/tapered capillaries in a packing bomb pressurized to 100 bar with N2. Sorbent suspension was prepared in methanol in a 2 ml flat-bottom glass vial (VWR Cat. No. 66020-950). The required quantity of the sorbent (see below) was added to the vial and mixed with methanol using a vortexer. Sorbent was left to swell for 30 min. and then vortexed and sonicated in an ultra-sonication bath for 60 s. The sorbent suspension vial was placed inside the pressure bomb. The capillary (fritted or tapered) was mounted with its open end (proximal end) dipped into the vial and fixed with a top nut. The pressure bomb system was pressurized to 100 bars by N_2_ gas. The capillary column was packed to a length that was 5 to 10 cm longer than the desired final column length. After packing, the pressure was slowly released over a 10 min period to avoid bubbling inside the column. The freshly packed capillary column was connected to and run on an LC system with 95% ACN at 150 nl/min flow rate (75 μm ID; for other IDs recalculate the flow rate proportionally to the capillary cross-section) for 30 min to compress the sorbent bed. The column was then cut to the desired length. Capillary columns were stored fully immersed in 10% aqueous ethanol.

### Standard packing (low sorbent concentration)

A volume of 400 μl low-concentration sorbent suspension (2-50 mg/ml) in methanol was prepared as described above. A magnet bar (2*3 mm) was put into the vial and rotated at 1000-1500 RPM to keep the sorbent evenly suspended during the packing process. The capillary was mounted in a pressure bomb with the open proximal end positioned 4-5 mm above the bottom of the vial with the sorbent suspension, without touching the rotating magnet. The rest of the protocol is as described above.

### *FlashPack* packing (high sorbent concentration)

Sorbent suspension preparation. 50-100 mg of sorbent (a 2-3 mm layer of dry sorbent in a flat-bottomed vial) was resuspended in 1 ml of methanol. The sorbent was left to settle by gravity for 20-30 min. The final settled sorbent particle layer on the bottom of the vial must be at least 5 mm deep. The vortex-sonication-settling cycle must be repeated if the sorbent suspension was not in use for more than 12 hours. A small magnet stirring bar (2 × 3 mm) was placed in the sorbent vial and set to rotate at a low speed (400-500 RPM).

**Capillary positioning**. An empty capillary (no solvent inside) was mounted in the packing bomb so that it reached more than 2 mm below the surface of the settled sorbent layer and 1-2 mm above the bottom of the vial. To achieve that the capillary was pushed all the way to the bottom of the vial, then retracted 1-2 mm and fixed with a nut. The capillary and the magnet bar must contact each other for the whole packing time, so the rotating magnet bar provides continuous mechanical “tapping” of the open proximal capillary end. The bomb was pressurized immediately after mounting the capillary in order to avoid passive filling of the capillary with the solvent.

The packing process was visually controlled. If the mechanical tapping of the proximal capillary end (cupola destabilization) was not effective, there was no appearance of opaque sorbent slurry being transferred into the capillary column and the capillary remained filled with the solvent and was fully transparent. In this case the magnet bar rotation speed was temporarily (3-4 s) increased to 1000 RPM. If the packing did not proceed, the pressure bomb was vented, and the capillary end repositioned. The remainder of the packing procedure was as described in the General Procedure above.

### Protein digestion

HeLa cell pellet or differentiating human myoblasts (- 80°C) were thawed at +4°C in solubilization buffer containing 50 mM TEAB, 1% SDC, 10 mM TCEP, 40 mM CAA, pH 8.5, supplemented with a cOmplete™ Protease Inhibitor Cocktail (Roche). Cells were lysed using an ultrasonic homogenizer Bioruptor^®^ in ice-water slurry. Protein concentration was measured by Trp fluorescence [14] and adjusted to 1 mg/ml with solubilization buffer. Samples were aliquoted into portions (100 μg of protein) and heated at 80°C for 10 min. Nine volumes of acetone (-20°C) was added and the samples were incubated overnight at -20°C and centrifuged at 15,000 g. The pellet was washed twice with acetone (-20°C) and air-dried for 10 min. 100 μl of 1% SDC in 50 mM TEAB buffer, pH 8.5 was added and the sample was heated for 5 min at 80°C to resolubilize the proteins. Protein digestion was carried out using 1 μg of trypsin per 100 μg of protein for 5 h at 37°C, subsequently an additional portion of trypsin (1 μg per 100 μg of protein) was added and the sample was incubated overnight at 37°C. The reaction was stopped using 10 μl of 10% TFA (final concentration ~1%). Precipitated SDC was removed by ethyl-acetate extraction [15, 16]. Peptide concentration was measured using Trp fluorescence [14]. Samples were stored frozen at -20°C. Prior LC-MS analysis, peptide samples were thawed and sonicated for 1 min and centrifuged for 15 min at 15,000 g.

### LC-MS

LC-MS analysis was carried out using an Ultimate 3000 RSLCnano HPLC system connected to a Fusion Lumos mass spectrometer (ThermoFisher Scientific). Capillary columns were conditioned using a BSA tryptic digest (50 fmol) in a 15 min LC gradient. Samples were loaded on PepMap 100 C18 5 μm 0.3 x 5 mm Trap Cartridge (ThermoScientific) in loading solvent (2% ACN, 98% H_2_O, 0.1% TFA) at 10 μl/min flow and separated with a linear gradient of 99.9% ACN, 0.1% FA (buffer B) in 99.9% H_2_O, 0.1% FA (solvent A) from 4 to 32% of solvent B in 2 h (for HeLa samples or myoblast samples) or from 2 to 25% B in 15 min (for BSA samples) at 0.25 μl/min flow. Separation was carried out either with a FlashPack columns packed with Inertsil sorbent, or with a commercially available EasySpray 75 μm* 50 cm PepMap 100 2μm column (ThermoFisher Scientific). Both the precolumn and the column were thermostated at 45°C (for 50 cm columns) or at 55°C (for 70 cm columns). An in-house developed micro-column-oven was used for *FlashPack* columns.

MS data was collected in DDA mode with 2 s cycle time. MS1 scans were collected in an Orbitrap mass-analyzer and MS2 scans were collected in a linear ion-trap. MS1 parameters were as follows: 120K resolution, 375-1500 scan range, max injection time 50 ms, AGC target 4×10^5^, RF Lens 30%. Ions were isolated with 1.6 m/z window targeting the highest intensity peaks of +2 to +7 charge and of 5x10^3^ minimum intensity. Dynamic exclusion was set to 30 s. MS2 fragmentation was carried out in CID mode with 35% CE. Ions were accumulated for max 300 ms with target AGC 3x10^3^. “Inject ions for all parallelizable time” was on. MS2 spectra were recorded in Rapid scan rate and saved in centroid datatype. ESI voltage was 2 kV for EasySpray columns, 2.5 kV for 50 cm and 2.9 kV for 70 cm FlashPack columns respectively.

### Data analysis

Raw spectra were processed using SkyLine (MacCossLab) [17] and FreeStyle (ThermoScientific). Peptide and protein identification was carried out using MaxQuant 1.5.7.0 (MQ) [18] and PEAKS 8.5 (Bioinformatics Solutions Inc)[19]. In both programs the data was searched against Swiss-Prot Homo Sapiens database (2016 Sep 29, 20211 entries).

*MaxQuant* was used for detailed LC peak analysis (retention time, peak width and FWHM) and for protein quantitation. Search was performed using the default MQ parameters. The following options were turned on: second peptide, maxLFQ, match between runs, calculate peak properties. All runs were analyzed as independent experiments. Reported identification numbers include only identified “by MS/MS” (i.e. not “by matching”). RT values (before alignment) and peak width (full peak width and FWHM) are derived from the result file allPeptides.txt.

*PEAKS* software was used for peptide peak area analysis using default PEAKS parameters and contaminant database (cRAP database, The Global Proteome Machine). Precursor mass tolerance was set to 5 ppm and fragment ion tolerance to 0.2 Da. Following parameters were used: trypsin [D | P]; max 2 missed cleavages; fixed Cys carbamidomethylation; default variable modifications: Deamidation (NQ), N-term Acetylation and Met Oxidation, with max 3 modification per peptide allowed. FDR estimation was carried out using the decoy-fusion method. Label free quantitation was performed only for protein database identifications (no *de novo)* with mass-error tolerance 20 ppm, RT shift tolerance 6 min and using TIC based normalization. All runs were marked as independent experiments. Peptides were selected with quality filter 2 and charge between 2 and 5. Protein filters were turned to minimum.

*Sample peak capacity, p_c_*, was calculated using the following formula

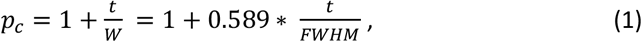
where *t*–full gradient time, *W*–average peak width at 4*σ* (13.5% peak height). Since there is no available proteomics software capable of direct *W* calculation for complex peptide samples with thousands of features, we used full width at half peak height *(FWHM)* instead, which is roughly equivalent to 2.356\**σ*.

*Chromatographic peak shape characteristics* (tailing factor, *T_f_*, and asymmetry factor, *A_s_*) were analyzed by manual EIC processing in a graphical software (Adobe PS) after EIC generation in FreeStyle (ThermoScientific). Normal distribution graph was generated in Excel and manually superimposed over EIC. *T_f_* and *A_s_* were calculated as

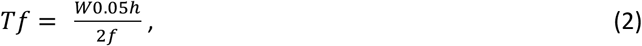

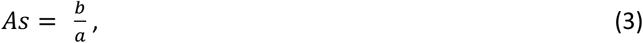

Where *W_0_._05h_* - full peak width at 5% of peak height, *f* – peak width at 5% of peak height to the left from peak apex time, *a* and *b* are peak widths to the left and to the right of the peak apex time respectively, measured at 10% of peak height.

Column quality and reproducibility assessments were performed based on the following parameters: overview of TIC/BPC profiles; retention time distribution; peak width (full peak width and FHWM); peak shape; number of identified peptides, number of identified proteins, reproducibility of peptide and protein quantitative measurements.

## Results and discussion

### Column entrance blocking in low pressure/high concentration packing

The goal of this study was to develop a technically simple, versatile low-pressure protocol for fast packing of long UHPLC capillary columns. To achieve this, it was necessary to overcome capillary entrance blocking by “sorbent clogging” during column packing at high sorbent concentrations (Supplementary Fig. S1a).

We rationalized that the blocking must be dynamic rather than static: instead of a large lump of sorbent aggregate blocking the entrance like a cork, there must be a dynamic self-assembling structure, probably resembling that of a brick cupola or dome in architecture [20] and not dissimilar to a self-assembling frit in a pulled emitter [6] (**Supplementary Fig. S2a-c**). The sorbent cupola-like structure will be stabilized by sorbent particle aggregation and solvent flow in the way mortar and gravity, respectively, stabilize a brick dome. Such self-assembling block/cupola is permeable for solvents, but it prevents additional sorbent material to enter into the capillary, effectively stopping column packing (**Fig. 1b**). Because the solvent flow is essential for the structure stability, the self-assembling cupola at the capillary entrance cannot be produced or kept outside the pressure bomb and thus cannot be directly visualized. However, a very similar blocking process takes place between two capillaries of diminishing internal diameters: while the larger ID capillary is being packed, the smaller ID capillary is closed to the sorbent by a self-assembling blocking structure (**Supplementary Fig. S2d**). We hypothesized that continuous destabilization of the cupola will allow packing at very high sorbent concentrations independently of system pressure.

### cUHPLC column packing by FlashPack method

Our *FlashPack* method combines mechanical block/cupola destabilization with column packing from ultra-high sorbent suspension concentration. Continuous cupola destabilization is conveyed by mechanical disturbance of the proximal capillary end by a small magnet bar. The capillary must be positioned deep enough in the sorbent slurry vial that the open end of the capillary touches the rotating magnet bar, which will continuously hit and tap it thus destabilizing and breaking any block/cupola of sorbent that interferes with the column packing (**Fig. 1c, Supplementary Fig. S3a & b**). Alternative destabilization approaches are also possible in the *FlashPack* method (**Supplemental Materials, Appendix A**). The column packing rate is proportional to the sorbent suspension concentration. The highest concentration can be achieved in the form of a gravity settled sorbent layer, where the concentration can go as high as 500-1000 mg/ml (**Supplemental Materials, Appendix B**).

The scheme of the packing process is shown on **Supplementary Fig. S4**. The capillary is mounted inside the vial with the preliminarily settled sorbent layer (**step 1**). Upon system pressurization the capillary gets filled with the concentrated sorbent slurry (**step 2**), while magnet bar rotation prevents block/cupola formation and stabilization and allows for unhindered continuous packing to the desired column length (**step 3**). The rotational speed of the magnet is kept to a minimum speed that is sufficient for cupola destabilization (400-500 RPM). High rotation speed leads to excessive sorbent grinding by a magnet bar [21] and resuspends the settled sorbent layer reducing the effective concentration and packing rate. When the destabilization works properly, the concentrated sorbent suspension moving inside the column can be followed with the naked eye as dense regions occasionally interrupted by transparent gaps (see **Supplementary Fig. S4**, **step 3**).

An empty capillary will get filled with a concentrated sorbent suspension even without intentional mechanical destabilization (**Supplementary Fig. S4, step 2**). We suggest that the very high flow rate can destabilize the cupola by itself in a way resembling a brick dome breaking under higher gravity. The effect might explain why column entrance blocking was never reported for high-pressure packing, which goes at proportionally higher flow rates than the low pressure packing. However, as the capillary gets filled with the liquid and the resin begins to get packed at the distal end of the column, the backpressure goes up, the flow rate drops and the cupola stabilizes. If no deliberate destabilization procedure is applied, no new sorbent can enter and the solvent flow packs up the sorbent already inside the column capillary (**Supplementary Fig. S4, step 4**). In effect, a short cHPLC column can be produced in a very short time with no additional mechanical destabilization needed. The amount of the sorbent getting inside the capillary during the filling step is proportional to the capillary length, so we recommend that the initial capillary length exceeds the desired column length by 50-100%. For example, a 50 cm capillary is recommended for packing 15-25 cm columns, and 80 cm capillaries for packing 50 cm columns using the *FlashPack* method.

The *FlashPack* protocol increases the packing speed and reduces the packing time by 100-fold (a few minutes) for standard capillary HPLC columns of 10-30 cm length. The time required to pack a 50 cm *FlashPack* cUHPLC column with 2 μm sorbent is less than one hour (**Fig. 1d & e**).

### Versatility of FlashPack method

The FlashPack packing procedure was tested with several types of RP sorbent, including classical fully porous silica sorbents (Inertsil, Reprosil, Luna 2, Zorbax, Luna Omega) as well as hybrid (BEH), polymeric (Triart) and core-shell (Aeris) sorbents, and HILIC and mixed-mode IEX-HILIC sorbents (polyLC and polyCAT) (data not shown). The size of the tested sorbent particles ranged from 1.6 to 5 μm. The *FlashPack* approach worked identically for all tested sorbents except for YMC Triart C18, which required less polar solvents, such as acetone or methanol/isopropanol mixture, for the sorbent suspension. Interestingly, initial filling of the capillary with the sorbent suspension (**Supplementary Fig. S4 steps 2, 4**) is not affected by the sorbent size, since the suspension displaces the air. Thus, short columns can be filled with 5 μm, 1.6 μm or future sub-1-μm sorbents in almost similarly short times.

We successfully tested the *FlashPack* protocol with fused silica capillaries of different inner diameters, from 30 to 200 μm (data not shown). *FlashPack* was effective for both fritted capillary columns and pulled (ESI taper-tip) capillary columns for direct interfacing to ESI MS. We noted that column packing at high sorbent slurry concentration improved the efficiency of self-organizing frit formation in the tapered column end, thereby reducing the rate of sporadic column blocking immediately after the bomb pressurization.

### FlashPack columns perform similarly to commercial columns

The columns produced with the *FlashPack* approach demonstrated excellent *separation performance, repeatability* and *durability*. A 50 cm *FlashPack* reversed-phase column was used over a two-month period for proteomics sample analysis for more than 200 sample injections. Comparative performance analysis to a commercial column with similar dimensions and sorbent characteristics was done before and after experimental samples (**Supplementary Fig. S5**). During the testing period, the *FlashPack* column was regularly disconnected and reconnected to the LC system. The *FlashPack* column demonstrated separation quality comparable to the commercial column both immediately after the packing and after extensive use without any noticeable degradation (**Fig. 2, Supplementary Fig. S5**). The column showed excellent run-to-run reproducibility (repeatability) and durability during a proteomics study of muscle cell (myoblast) differentiation (**Supplementary Fig. S6**). The quantitative proteomics data obtained using *FlashPack* columns was in accordance with previously published results [22]. Our results reconfirmed that column packing from high sorbent slurry concentrations did not negatively affect chromatographic resolution [12, 21, 23].

**Fig. 2.**
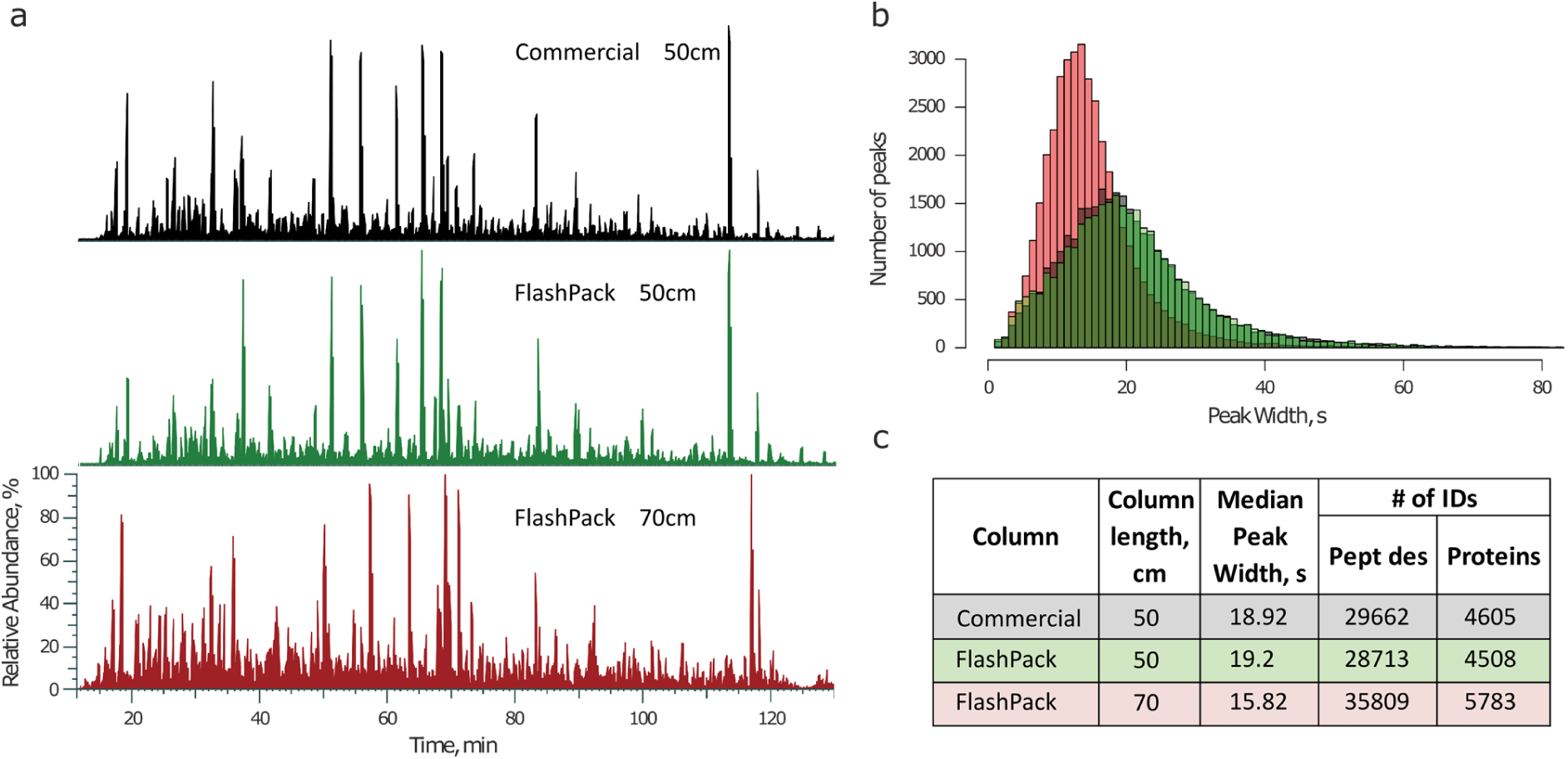
Comparison of commercial (50 cm) and *FlashPack* columns (50 and 70 cm) performance in a 2 hour LC-MS analysis of a HeLa cell tryptic digest. **(a)** Alignment of base peak chromatograms of a commercial column (black trace),50 cm *FlashPack* column (green trace) and 70 cm *FlashPack* column (red trace) show close similarity in peak features and separation efficiency between commercial and FlashPack columns. **(b)** Baseline peak width distribution for all unique identified peptides is almost identical between 50 cm commercial (black) and FlashPack (green) columns (see Supplementary Fig. S5 for more details). A 70 cm *FlashPack* column (red) provides improved chromatographic resolution. **(c)** Performance of the 50 cm *FlashPack* column is very similar to the commercial column of identical dimensions (single LC-MS run data is shown). The longer 70 cm FlashPack column gives sharper chromatographic peaks and more peptide and protein identifications in the same analysis time (an average result for 5 independently packed columns).

### FlashPack column packing is highly reproducible

To demonstrate the reproducibility and analytical performance of *FlashPack* column based LC-MS, a HeLa cell tryptic digest was analyzed using five 70 cm long and 75 μm ID *FlashPack* ESI emitter columns (**Fig. 3**). The chromatographic resolution increase between 50 and 70 cm capillary column lengths corresponded to the expected 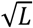 proportionality (eq. 5, [24]). The median full peak width was on average improved from 19 to 16 s and the number of peptide IDs per LC-MS run increased from 29,000 unique peptides to > 35,000 peptides originating from 5780 proteins (**Supplementary Fig. S7**). The FWHM standard deviation between the five tested columns was 8.67%, which was comparable to that reported for ultra-high pressure packing protocols [25].

**Fig. 3.**
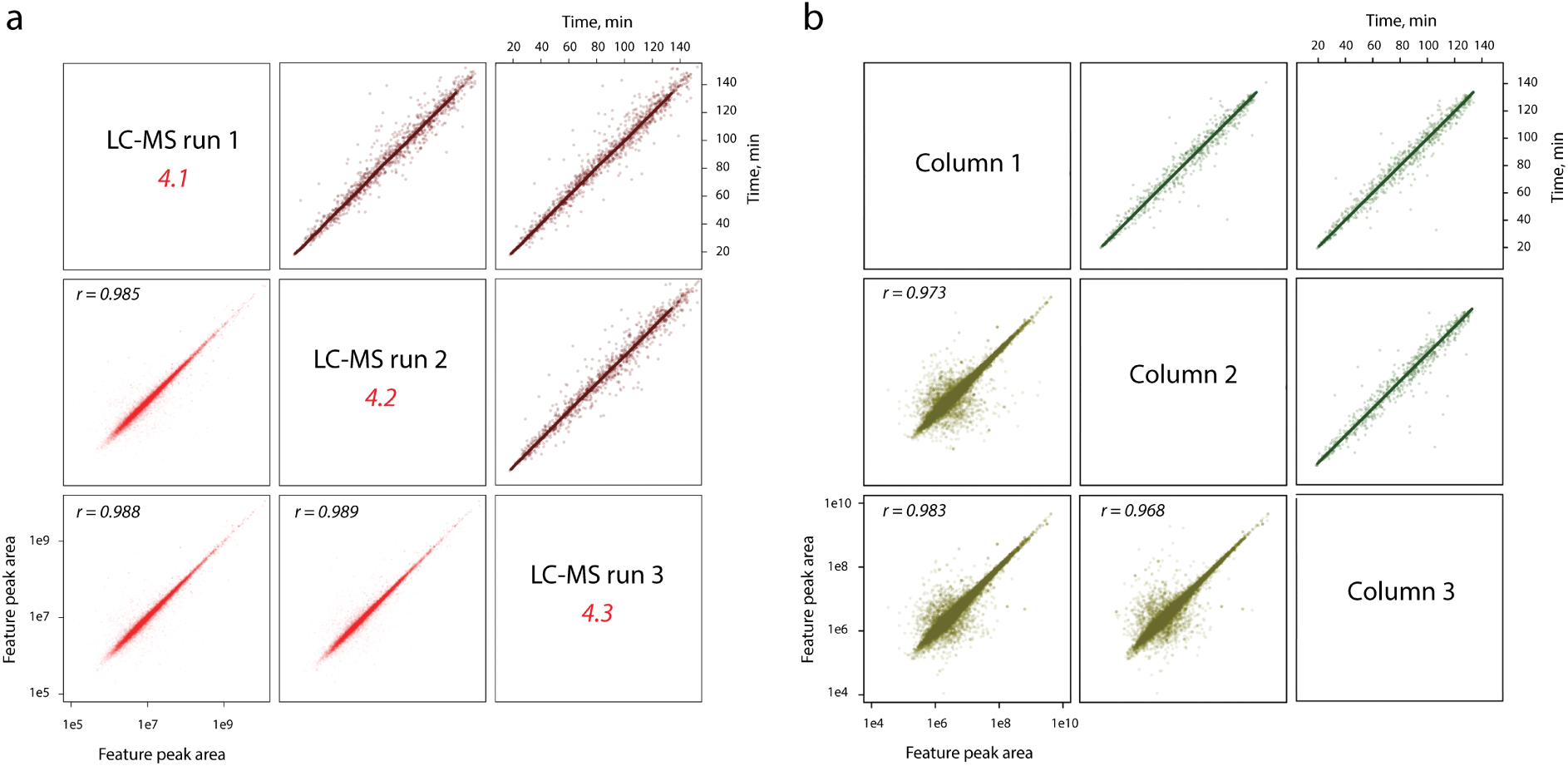
The *FlashPack* method produces columns with stable separation properties over several runs and between independently prepared columns. **(a)** Comparison of 3 repeat LC-MS runs using a 50 cm *FlashPack* column, human myocyte proteome sample and 2h gradient shows high correlation of retention time (brown trace) and quantitative peptide measurement (red trace); **(b)** Comparison of 3 independently prepared 70 cm FlashPack columns tested using HeLa cells extract and 2h LC-MS analysis; (dark green trace) - retention time; (light green trace) - peptide quantitation. The complete analysis is provided in Supplementary Fig. S6 and S7.

### Conclusions

The *FlashPack* approach produces high quality cUHPLC columns for bioanalytical applications. It enables interested researchers to pursue experiments with ultra-high performance column chromatography using various sorbents and column size for applications of cUHPLC and LC-MS in biology and biomedicine. The *FlashPack* method uses a standard pressure bomb that is already present in many LC-MS laboratories and thus can be quickly implemented.

## Author contributions

S.K. designed and executed experiments; S.K., O.N.J., and A. R. contributed to data analysis and interpretation, and wrote the manuscript.

## Acknowledgements

We thank Frank Mortensen and Torben Christensen from the SDU Department of Biochemistry and Molecular Biology workshop for manufacturing the micro-column oven used in this study. SK and ARW were supported by a grant from the Independent Research Fund Denmark - Natural Sciences (to ONJ). We thank the VILLUM Foundation for a generous grant to the VILLUM Center for Bioanalytical Sciences at SDU (to ONJ).

## Data availability

The mass spectrometry proteomics data is deposited at the ProteomeXchange Consortium (http://proteomecentral.proteomexchange.org) via the PRIDE partner repository [26] with the dataset identifier PXD009519.Figure legends

## Abbreviations

ACN: acetonitrile
BSA: bovine serum albumin
CAA: 2-Chloroacetamide
cUHPLC: capillary ultra-high performance liquid chromatography
DB: database
EIC: extracted ion current chromatogram
ESI: electro-spray ionization
FA: formic acid
FDR: false discovery rate
FS: fused silica
FW: full width
FWHM: full width at half maximum
ID: internal diameter
LC: liquid chromatography
LC-MS: liquid chromatography-mass spectrometry
MQ: MaxQuant
OD: outer diameter
RT: retention time
SDC: Sodium deoxycholate
TCEP: Tris(2-carboxyethyl)phosphine
TEAB: triethylammonium bicarbonate
TFA: trifluoric acid
TIC: total ion current chromatogram

## References

1. Bekker-Jensen, D.B., et al., An Optimized Shotgun Strategy for the Rapid Generation of Comprehensive Human Proteomes. Cell Syst, 2017. 4(6): p. 587–599 e4.

2. Hebert, A.S., et al., The one hour yeast proteome. Mol Cell Proteomics, 2014. 13(1): p. 339–47.

3. Shishkova, E., A.S. Hebert, and J.J. Coon, Now, More Than Ever, Proteomics Needs Better Chromatography. Cell Syst, 2016. 3(4): p. 321–324.

4. Detailed explanation on the workings of the pressure bomb packing setup. https://www.nextadvance.com/pressure-injection-cells-lc-ms-capillary-column-packing-loader/?target=Overview. Available from: https://www.nextadvance.com/pressure-injection-cells-lc-ms-capillary-column-packing-loader/?target=Overview.

5. Cortes, H.J., et al., Porous ceramic bed supports for fused silica packed capillary columns used in liquid chromatography. Journal of High Resolution Chromatography, 1987. 10(8): p. 446–448.

6. Ishihama, Y., et al., Microcolumns with self-assembled particle frits for proteomics. J Chromatogr A, 2002. 979(1–2): p. 233–9.

7. Yates, J.R., 3rd, et al., Future prospects for the analysis of complex biological systems using micro-column liquid chromatography-electrospray tandem mass spectrometry. Analyst, 1996. 121(7): p. 65r–76r.

8. Wang, F., et al., Integration of monolithic frit into the particulate capillary (IMFPC) column in shotgun proteome analysis. Anal Chim Acta, 2009. 652(1–2): p. 324–30.

9. Liu, H., et al., Effects of column length, particle size, gradient length and flow rate on peak capacity of nano-scale liquid chromatography for peptide separations. J Chromatogr A, 2007. 1147(1): p. 30–6.

10. MacNair, J.E., K.C. Lewis, and J.W. Jorgenson, Ultrahigh-pressure reversed-phase liquid chromatography in packed capillary columns. Anal Chem, 1997. 69(6): p. 983–9.

11. Jorgenson, J.W., Capillary liquid chromatography at ultrahigh pressures. Annu Rev Anal Chem (Palo Alto Calif), 2010. 3: p. 129–50.

12. Bruns, S., et al., Slurry concentration effects on the bed morphology and separation efficiency of capillaries packed with sub-2 mum particles. J Chromatogr A, 2013. 1318: p. 189–97.

13. Maiolica, A., D. Borsotti, and J. Rappsilber, Self-made frits for nanoscale columns in proteomics. Proteomics, 2005. 5(15): p. 3847–50.

14. Wisniewski, J.R. and F.Z. Gaugaz, Fast and sensitive total protein and Peptide assays for proteomic analysis. Anal Chem, 2015. 87(8): p. 4110–6.

15. Masuda, T., M. Tomita, and Y. Ishihama, Phase transfer surfactant-aided trypsin digestion for membrane proteome analysis. J Proteome Res, 2008. 7(2): p. 731–40.

16. Leon, I.R., et al., Quantitative assessment of in-solution digestion efficiency identifies optimal protocols for unbiased protein analysis. Mol Cell Proteomics, 2013. 12(10): p. 2992–3005.

17. MacLean, B., et al., Skyline: an open source document editor for creating and analyzing targeted proteomics experiments. Bioinformatics, 2010. 26(7): p. 966–8.

18. Cox, J. and M. Mann, MaxQuant enables high peptide identification rates, individualized p.p.b.-range mass accuracies and proteome-wide protein quantification. Nat Biotechnol, 2008. 26(12): p. 1367–72.

19. Zhang, J., et al., PEAKS DB: de novo sequencing assisted database search for sensitive and accurate peptide identification. Mol Cell Proteomics, 2012. 11(4): p. M111 010587.

20. Dahmen, J.F.D. and J.A. Ochsendorfs, 17 - Earth masonry structures: arches, vaults and domes, in Modern Earth Buildings, M.R. Hall, R. Lindsay, and M. Krayenhoff, Editors. 2012, Woodhead Publishing. p. 427–460.

21. Wahab, M.F., et al., Fundamental and Practical Insights on the Packing of Modern High-Efficiency Analytical and Capillary Columns. Anal Chem, 2017. 89(16): p. 8177–8191.

22. Le Bihan, M.C., et al., Cellular Proteome Dynamics during Differentiation of Human Primary Myoblasts. J Proteome Res, 2015. 14(8): p. 3348–61.

23. Reising, A.E., et al., Bed morphological features associated with an optimal slurry concentration for reproducible preparation of efficient capillary ultrahigh pressure liquid chromatography columns. J Chromatogr A, 2017. 1504: p. 71–82.

24. Gilar, M., et al., Implications of column peak capacity on the separation of complex peptide mixtures in single- and two-dimensional high-performance liquid chromatography. J Chromatogr A, 2004. 1061(2): p. 183–92.

25. Godinho, J.M., et al., Implementation of high slurry concentration and sonication to pack high-efficiency, meter-long capillary ultrahigh pressure liquid chromatography columns. J Chromatogr A, 2016. 1462: p. 165–9.

26. Vizcaino, J.A., et al., 2016 update of the PRIDE database and its related tools. Nucleic Acids Res, 2016. 44(D1): p. D447–56.

